# SK609, a novel dopamine D3 receptor agonist and norepinephrine transporter blocker with pro-cognitive actions, does not induce psychostimulant-like increases in risky choice during probabilistic discounting

**DOI:** 10.1101/2024.06.05.597571

**Authors:** Christopher P. Knapp, Brooke Fallon, Sandhya Kortagere, Barry D. Waterhouse, Stan B. Floresco, Rachel L. Navarra

**Affiliations:** Department of Cell Biology and Neuroscience, Rowan-Virtua School of Translational Biomedical Engineering and Sciences, School of Osteopathic Medicine, Stratford, NJ, USA; Department of Microbiology and Immunology, Drexel University, Philadelphia, PA, USA; Department of Psychology and Djavad Mowafaghian Centre for Brain Health, University of British Columbia, Vancouver, BC, Canada

**Keywords:** Amphetamine, Methylphenidate, Norepinephrine Transporter, Dopamine D3 Receptor, Decision Making

## Abstract

**Rationale:** Psychostimulants, such as amphetamine (AMPH) and methylphenidate (MPH), non-selectively elevate extracellular concentrations of the catecholamine neurotransmitters, dopamine (DA) and norepinephrine (NE), and are common pharmacological strategies used to improve prefrontal cortex (PFC)-dependent cognitive dysfunction. However, this approach can be problematic given AMPH has been shown to increase preference for risky choices in a rodent assay of risk/reward decision making. SK609 is a novel NE reuptake blocker that selectively activates DA D3 receptors without affinity for the DA transporter. SK609 has been shown to improve cognitive performance without increasing psychostimulant-like spontaneous locomotor activity, suggesting SK609 may benefit neurocognitive function without psychostimulant-like side effect liability.

**Objectives:** We compared AMPH, MPH, and SK609 within dose ranges that display their cognitive enhancing properties in a probabilistic discounting task (PDT) of risk/reward decision making behavior to assess their potential to increase risky choice preference.

**Methods:** Rats chose between small/certain rewards delivered with 100% certainty and large/risky rewards delivered with descending probabilities across a session (100-6.25%) following administration of AMPH (0.25-1 mg/kg), MPH (2-8 mg/kg), and SK609 (4 mg/kg).

**Results:** AMPH and MPH increased risky choice behavior at doses previously reported to enhance cognition, whereas SK609 did not. AMPH and MPH also reduced sensitivity to non-rewarded risky choices.

**Conclusions:** These data highlight the combination of NE transporter blockade and selective D3 activation in pro-cognitive action without psychostimulant-like side effect liability. The absence of DA transporter blockade and non-selective dopaminergic activation are beneficial properties of SK609 that differentiates it from the traditional pro-cognitive psychostimulants.

## Introduction

Cognitive and reward-related processes that drive decision making during goal-directed behaviors are modulated by catecholamine inputs to the corticolimbic system, such as dopamine (DA) and norepinephrine (NE) projections to the prefrontal cortex (PFC) (Arnsten, 1998; Arnsten and Li, 2005; Robbins and Arnsten, 2009; Logue and Gould, 2014). Decision making is a supraordinate executive function that requires integration of flexibility, action/outcome monitoring, and cost/benefit estimations. Optimal operation of these processes is critical in choosing beneficial actions, particularly in situations involving risk or uncertainly (Shead and Hodgins, 2009). Impaired decision making has been observed in clinical populations with neurological deficits that have been linked to imbalanced catecholamine function within the PFC, such as attention deficit-hyperactivity disorder (ADHD), substance use disorders, and following traumatic brain injury (TBI) (Buelow and Suhr, 2009).

The Iowa Gambling Task (IGT) is the most widely used clinical test to assess risk/reward decision making in humans. The IGT requires subjects to make choices that involve uncertain opportunities to win money (Bechara et al., 1994). Subjects must choose cards from decks associated with different schedules of reinforcement-punishment contingencies, e.g. ‘safe’ options that give small rewards but also result in small penalties and ‘risky’ options that give large rewards but result in large penalties. As such, participants must keep track of choice/outcome history to adjust decision biases accordingly. Risk/reward decision making can be evaluated in pre-clinical rodent models using probabilistic discounting tasks (PDT) (St Onge and Floresco, 2009). The PDT requires rats to choose between a small/certain reward that is delivered with 100% certainty or a large/risky reward that is delivered with decreasing probabilities (100, 50, 25, 12.5, and 6.25%) over a test session. Normal animals show a characteristic “probabilistic discounting curve” as the probability of reward decreases, demonstrating that the subjective value of the larger reward decreases with the probability of receiving it. The PDT has previously been used to evaluate pharmacological treatments that augment or reduce DA and NE activity to reveal the specific mechanisms in which catecholaminergic circuits mediate risk/reward decision making (Floresco and Whelan, 2009; Montes et al., 2015; St Onge and Floresco, 2009; St Onge et al., 2010; Stopper et al., 2013).

Psychostimulant drugs block reuptake and elevate extracellular concentrations of both DA and NE, and are common pharmacological strategies used to improve PFC-mediated executive dysfunction associated with ADHD (De Crescenzo et al., 2017) and more recently following TBI (Yamamoto et al., 2018; LeBlond et al., 2019; Kurowski et al., 2019; Ekinci et al., 2017; Al-Adawi et al., 2020; Huang et al., 2016; DeMarchi et al., 2005; Levin et al., 2019; Traeger et al., 2020; Johansson et al., 2017; Warden et al., 2006; Plenger et al., 1996). However, this approach can be problematic given adverse psychostimulant-associated side effect profiles and their potential to further increase risk seeking behaviors. For example, the psychostimulant drug amphetamine (AMPH) induces less advantageous decision making behavior by increasing preference for risky choices in healthy humans performing the IGT (Oswald et al., 2015) as well as rodents performing the PDT (St Onge et al., 2010; St Onge and Floresco, 2009). Thus, by blocking DA and NE transporters and enhancing DA release with AMPH, preferences for larger, probabilistic rewards increase even as their long-term value decreases.

Additional reports have further characterized drugs that target specific catecholamine transporters and receptor systems that influence decision making using the PDT. Systemic activation of dopamine D1 and D2 receptors (St Onge and Floresco, 2009) and blockade of noradrenergic α2 receptors (Montes et al., 2015) increases risky choice when the likelihood of obtaining a probabilistic reward decreases over time. Additionally, blocking NET alone also increased risky choice preference, but only at high probabilities where this option was advantageous (Montes et al., 2015). In contrast, systemic antagonism of D1 and D2 receptors (St Onge et al., 2010; St Onge and Floresco, 2009) or stimulating the D3 or α2 receptors, reduces risky choice in the PDT (with the latter effects potentially driven by autoreceptor activation) (Montes et al., 2015; St Onge and Floresco, 2009). Therefore, reducing activity at certain catecholamine receptors can reduce the allure of larger yet riskier rewards. Together, these studies have established a framework for how catecholamine-based treatments can alter probabilistic discounting, and as such, this assay may be useful for testing novel, potential pro-cognitive compounds targeting these systems.

SK609 is a novel small molecule drug that blocks reuptake of NE and selectively activates DA D3 receptors without affinity for the DA transporter (Xu et al., 2017; Marshall et al., 2019). SK609 has been shown to improve sustained attention performance similar to the ADHD-approved psychostimulant methylphenidate (MPH, which also blocks DA and NE transporters), but not produce MPH-induced increases in spontaneous locomotor activity associated with blocking DA transporter activity (Marshall et al., 2019). These findings suggest SK609 may help ameliorate neurocognitive impairments without psychostimulant-like motor side effects. Of particular interest, these authors showed that SK609 more prominently improved this dimension of cognition in low-performing animals as compared to high-performing animals, suggesting SK609 may produce beneficial effects for animals that exhibit a range of neurocognitive impairments in behavior. In the present set of experiments, we evaluated AMPH, MPH, and

SK609 in the PDT to compare these three agents with cognitive-enhancing properties for their potential to increase risky choice behavior as a side effect.

## Materials and Methods

### Animals

Thirty male Long-Evans rats were used in this study. Animals were obtained at 225-250g from Charles River Laboratories and housed in a 12h:12h reverse light/dark cycle facility. Following a week of acclimation, rats were single housed into separate cages, and placed on a food regulated diet (5g per 100g body weight/ day) with *ad libitum* access to water. They were maintained to 85% of their free feeding weight throughout the duration of these studies. All experimental procedures were in accordance with the Rowan University School of Osteopathic Medicine Institutional Animal Care and Use Committee and the National Institutes of Health Guide for the Care and Use of Laboratory Animals.

### Probabilistic Discounting Task

#### Apparatus

Behavioral studies were conducted in 16 operant chambers [29cm (L) x 24cm (W) x 29cm (H); Med-Associates, Albans, VT] enclosed within sound attenuating boxes. Operant chambers were equipped with a fan, a house light, and 2 retractable levers located on either side of a food dispenser where sucrose pellet rewards (45 mg; Bio-Serv, Flemington, NJ) were delivered. A photo beam was located at the dispenser entry point to detect reward collection. Custom built Med Associates Nexlink computer packages controlled the training, testing, and data acquisition during performance.

#### Lever-Pressing Training

Initial training protocols were the same as those described previously (Knapp et al., 2024), which were adapted from St. Onge and Floresco (St Onge and Floresco, 2009). Rats were first trained to press a single lever (either the left or right) using a fixed-ratio one (FR1) schedule to a criterion of 50 presses within 30 minutes. Once criterion was achieved, rats repeated this procedure for the opposite lever. Rats then trained on a simplified version of the PDT (90 trials per session) which required them to press one of the two levers within a 10 second period for a sucrose reward delivered with a 50% probability. This procedure familiarized them with the probabilistic nature of actions and outcomes. Rats were trained for at least 3 days to a criterion of 75 or more successful trials (i.e., ≤ 15 omissions) on the simplified PDT.

#### PDT Training and Testing

The PDT was used to assess changes in risk/reward decision making and has been described previously (Floresco and Whelan, 2009; Montes et al., 2015; St Onge et al., 2011; St Onge et al., 2010; St Onge and Floresco, 2009; St Onge and Floresco, 2010; Stopper et al., 2014; Bercovici et al., 2023; Knapp et al., 2024). This task required rats to choose between levers that result in either small/certain rewards (1 pellet) delivered with 100% certainty and large/risky rewards (4 pellets) delivered with decreasing probabilities across a series of five trial blocks (i.e. 100% probability → 50% → 25% → 12.5% → 6.25%). Each session took 52.5 minutes to complete and consisted of 90 trials, separated into 5 blocks of 18 trials. These 18 trials consisted of 8 forced-choice trials where only one lever was extended allowing rats to learn the relative likelihood of obtaining the larger reward in each block. This was followed by 10 free-choice trials, where both levers were extended allowing rats to freely choose between the small/certain or the large/risky lever. Each session began in darkness with both levers retracted. A trial began every 35 seconds with the illumination of the house light and extension of one or both levers. Once a lever was chosen, both levers retracted, rats were rewarded 1 pellet if they chose the small/certain lever or a possible 4 pellets if they chose the large/risky lever, and the house lights turned off. If the rat did not respond within a 10 second period, levers retracted, and the house light turned off until the next trial and the trial was scored as an omission.

Rats were trained 7 days per week until each cohort achieved baseline criteria, which included choosing the risky lever in >80% of trials in the 100% block and maintaining stable patterns of choice for 3 consecutive sessions.

Determining stable baseline performance involved analyzing 3 consecutive sessions using a repeated measures analysis of variance (ANOVA) with two within-subjects factors (day and trial block). Rats were required to demonstrate a significant main effect of block (p < 0.05), but not a main effect of day nor a day x block interaction (p > 0.1). If animals, as a group, met these 3 requirements, they were determined to have achieved stable baseline levels of choice behavior.

Rats required ~18 days of PDT training before stable criterion performance was achieved. They then received approximately 10 additional days of training to maintain stable performance. A counter-balanced drug dosing design was then used to assign an order of drug or vehicle (saline) tests to each rat. This method was used to limit bias when forming surgical groups by having equal amounts of risky and risk-adverse rats in each group.

### Drug Testing

A within-subjects design was used for all drug testing with AMPH (Sigma-Aldrich), MPH (Sigma-Aldrich), and SK609 (PolyCore Therapeutics). All drugs were dissolved in sterile saline and injected intraperitoneally (i.p.) at a volume of 1 mL/kg either 10 min (AMPH), 15 min (MPH) or 5 min (SK609) before testing. AMPH was previously shown to increase risky choice in the PDT (St Onge and Floresco, 2009; Floresco and Whelan, 2009; St Onge et al., 2010). In the present experiments, we evaluated AMPH at a low dose (0.25 mg/kg) that has been shown to improve cognitive (Andrzejewski et al., 2014; Berridge and Stalnaker, 2002), a moderate dose (0.5 mg/kg) (Grilly and Loveland, 2001), and a high dose (1 mg/kg) above the drug’s therapeutic window that has been shown to promote hyperactive and impulsive behavior in rats (Dalia and Wallace, 1995; Grilly and Loveland, 2001). We evaluated the effects of MPH using a low dose (2 mg/kg) based within the range that has been shown to enhance cognitive performance or improve impulsive behavior in other tasks (Navarra et al., 2008; Jentsch et al., 2009; Paterson et al., 2011; Bizarro et al., 2004; Robinson, 2012; Berridge et al., 2012; Seu et al., 2009; van Gaalen et al., 2006; Navarra et al., 2017), as well as a high dose (8 mg/kg) that exceeds the therapeutic window and that we suspected would promote hyperactivity and impulsivity (Amini et al., 2004; Gaytan et al., 1996) and increase risk taking behavior. We then assessed SK609 at its cognitive enhancing dose (4 mg/kg) (Marshall et al., 2019). A counter-balanced drug dosing design was used to assign rats to an order of drug, dose, and vehicle for each test day. All rats received all doses of AMPH (n = 30), and then were split into two separate groups for testing with either MPH (n = 15) or SK609 (n = 12). Following drug test days, rats were retrained until they, as a group, displayed stable patterns of choice for 3 consecutive sessions, after which subsequent drug tests were administered. These 3 days also served as a washout period between drug doses. This procedure was repeated until all rats had received each of the designated treatments. Rats that did not return to stable baseline within any washout periods in which the majority met criteria (n = 3) were removed from subsequent testing.

### Data Analysis

Choice behavior was defined as the percent choice of the large/risky option, where the number of large/risky lever presses was divided by the total number of successful trials for each block of free-choice trials. Choice latency was recorded as the time elapsed between extension of both levers and a choice response. Magazine latency was recorded as the time elapsed between a choice response and head entry into the reward magazine, which was detected by breakage of the infrared beam. Drug-induced alterations in sensitivity to reward and negative feedback were assessed by win-stay/lose-shift analysis, which evaluated each response according to the outcome of the preceding trial (Stopper et al., 2013). A win-stay response was demonstrated when a rat selected the large/risky option on a trial immediately after winning a large/risky reward, whereas a lose-shift response was scored when the rat chose the small/safe option after a non-rewarded large/risky choice. The number of win-stay and lose-shift responses were then divided by the number of free-choice responses that resulted in wins or losses, respectively, across all trial blocks and expressed as a ratio.

All statistical analyses were performed using GraphPad Prism software (GraphPad Software, San Diego CA). Choice behavior, choice latency, and magazine latency were analyzed using two-way mixed-design ANOVAs with trial block (100%, 50%, 25%, 12.5%, and 6.25%) and treatment (AMPH – 0, 0.25, 0.5, and 1mg/kg; MPH – 0, 2, and 8 mg/kg; SK609 – 0 and 4 mg/kg) as within-subjects factors. In all of these analyses, the main effect of block was always significant (p < 0.001) for choice behavior and will not be discussed further. Similarly, Win-Stay/Lose-Shift behavior was analyzed using two-way repeated measures ANOVAs with feedback (win-stay and lose-shift) and treatment as within-subjects factors. Trial Omissions were analyzed using either two-way repeated measures or mixed-design ANOVAs with trial block (100%, 50%, 25%, 12.5%, and 6.25%) and treatment (AMPH – 0, 0.25, 0.5, and 1mg/kg; MPH – 0, 2, and 8 mg/kg; SK609 – 0 and 4 mg/kg) as within-subjects factors. Dunnett’s or Sidak’s multiple comparisons tests, when appropriate, were used to compare individual differences when overall significance was found. For all results, statistical significance was determined by a p value < 0.05.

## Results

### Effects of Amphetamine

Analysis of choice data (**Fig. 1a**) revealed a significant main effect treatment [F (2.437, 70.68) = 58.28, p < 0.0001] as well as a significant block x treatment interaction [F (5.181, 148.9) = 20.84, p < 0.0001]. Dunnett’s multiple comparisons analysis determined that rats treated with all doses of AMPH displayed a significant increase in risky choice (p < 0.05) in the 50%, 25%, and 12.5% blocks in comparison to vehicle (saline), whereas only the 0.5 and 1.0 mg/kg doses resulted in a significant increase in risky choice (p < 0.05) in the 6.25% block.

**Fig. 1.**
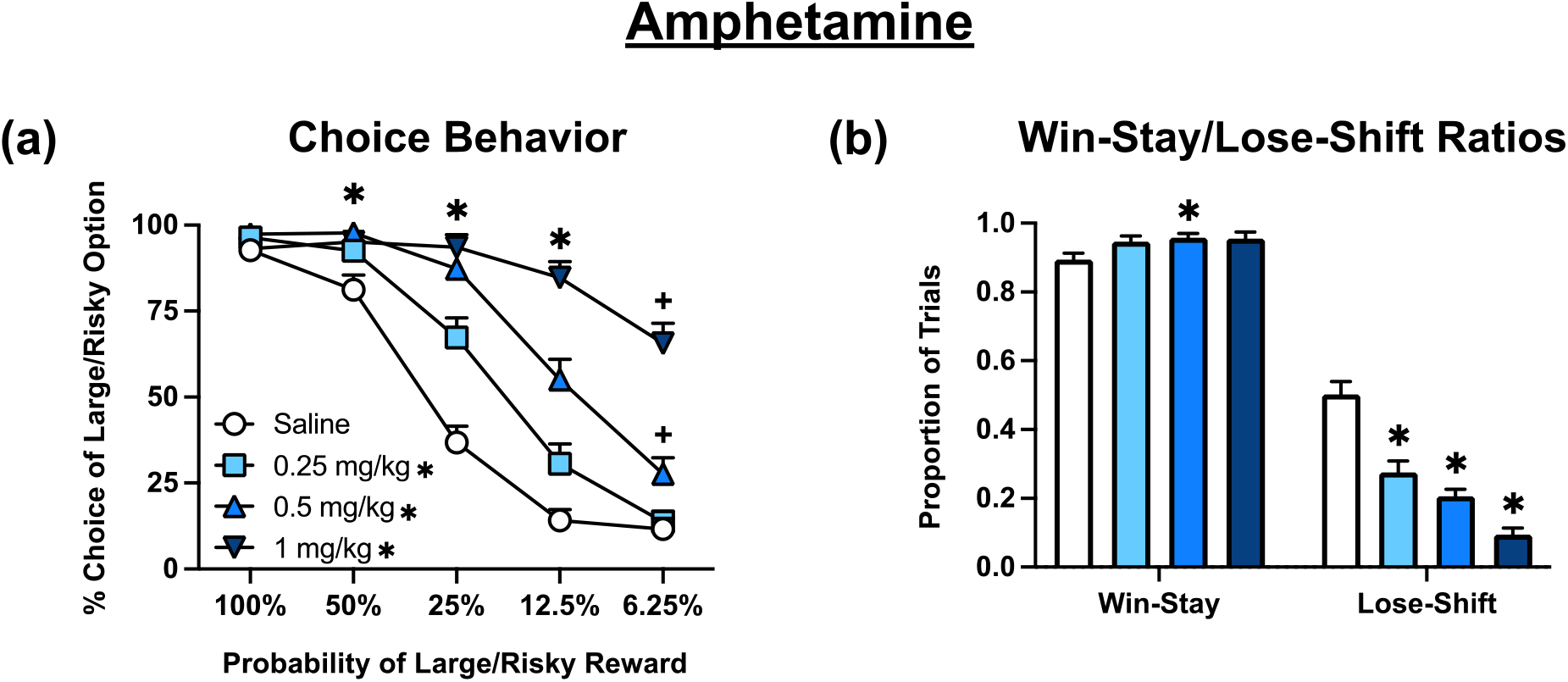
Effects of amphetamine (n = 30) during probabilistic discounting. (**a)** Choice Behavior: the percentage choice for the large/risky behavior over free choice trials is plotted across large/risky lever probability by block. * on symbol key denotes p < 0.05 for an overall treatment effect of each dose versus vehicle. * on graph denotes p < 0.05 for the 0.25, 0.5, and 1 mg/kg doses versus vehicle and + denotes p < 0.05 for the only the 0.5 and 1 mg/kg doses versus vehicle at a specific block analyzed by Dunnett’s multiple comparisons tests. (**b)** Win-Stay/Lose-Shift Ratios: the number of win-stay and lose-shift responses divided by the number of free-choice responses that resulted in wins or losses, respectively, across all trial blocks and expressed as a ratio. * on graph denotes p < 0.05 versus vehicle analyzed by Dunnett’s multiple comparisons tests. Symbols and bars represent mean ± SEM.

Analysis of win-stay/lose-shift data (**Fig. 1b**) revealed a significant main effect of both feedback [F (1.000, 29.00) = 914.2, p < 0.0001], and treatment [F (2.501, 72.54) = 21.63, p < 0.0001] as well as a significant feedback x treatment interaction [F (2.469, 71.61) = 36.34, p < 0.0001]. Dunnett’s multiple comparisons analysis determined that rats treated with all doses of AMPH displayed a significant decrease in lose-shift tendencies (p < 0.001) compared to vehicle-treated rats, whereas the 0.5 dose also resulted in an increase in win-stay behavior (p < 0.05).

Analysis of both choice and magazine latency data as well as trial omissions (**Table 1**) revealed a significant main effect of block [F (2.302, 66.75) = 6.501, p = 0.0017; F (2.646, 76.73) = 5.736, p = 0.0021; F (1.873, 54.33) = 6.407, p = 0.0038, respectively], but did not yield a significant main effect of treatment [F (2.347, 68.06) = 0.3345, p = 0.7505; F (2.273, 65.91) = 0.7875, p = 0.4738; F (1.291, 37.43) = 1.371, p = 0.2578, respectively] or a block x treatment interaction [F (5.836, 167.8) = 2.000, p = 0.0703; F (2.953, 79.72) = 2.551, p = 0.0624; F (2.538, 73.61) = 0.2939, p = 0.2939, respectively].

**Table 1.** Effects of Amphetamine, Methylphenidate, and SK609 on Choice Latency, Magazine Latency, and Trial Omissions.

| Drug Test | Choice Latency (sec) | Magazine Latency (sec) | Omissions |
| --- | --- | --- | --- |
| **Amphetamine** |  |  |  |
| Vehicle (saline) | 0.71 ± 0.09 | 0.40 ± 0.07 | 0.27 ± 0.12 |
| 0.25 mg/kg | 0.65 ± 0.07 | 0.44 ± 0.10 | 0.20 ± 0.10 |
| 0.5 mg/kg | 0.73 ± 0.09 | 0.49 ± 0.12 | 0.21 ± 0.09 |
| 1 mg/kg | 0.72 ± 0.09 | 0.57 ± 0.20 | 0.57 ± 0.28 |
| **Methylphenidate** |  |  |  |
| Vehicle (saline) | 0.70 ± 0.25 | 0.67 ± 0.12 | 0.45 ± 0.41 |
| 2 mg/kg | 0.67 ± 0.12 | 0.53 ± 0.15 | 0.21 ± 0.15 |
| 8 mg/kg | 1.02 ± 0.22 | 0.67 ± 0.24 | 0.96 ± 0.51 |
| **SK609** |  |  |  |
| Vehicle (saline) | 0.96 ± 0.18 | 0.26 ± 0.03 | 0.60 ± 0.28 |
| 4 mg/kg | 0.97 ± 0.21 | 0.20 ± 0.02 | 0.33 ± 0.17 |

Rats were administered amphetamine (AMPH – 0, 0.25, 0.5, and 1 mg/kg, n = 30), methylphenidate (MPH – 0, 2, and 8 mg/kg, n = 15), and SK609 (0 and 4 mg/kg, n = 12), and then tested in the probabilistic discounting task (PDT). No differences in choice latencies, magazine latencies, or trial omissions were observed. Values represent mean ± SEM averaged across blocks.

### Effects of Methylphenidate

Analysis of choice data (**Fig. 2a**) again revealed a significant main effect treatment [F (1.491, 20.88) = 20.12, p < 0.0001], that reflected an overall increase in risky choice induced by both doses (p < 0.05). The analysis also yielded block x treatment interaction [F (4.217, 57.99) = 14.79, p < 0.0001]. Dunnett’s multiple comparisons analysis determined that rats treated with the 8 mg/kg dose of MPH displayed a significant increase in risky choice preference (p < 0.05) in the 25%, 12.5%, and 6.25% blocks in comparison to vehicle (saline).

**Fig. 2.**
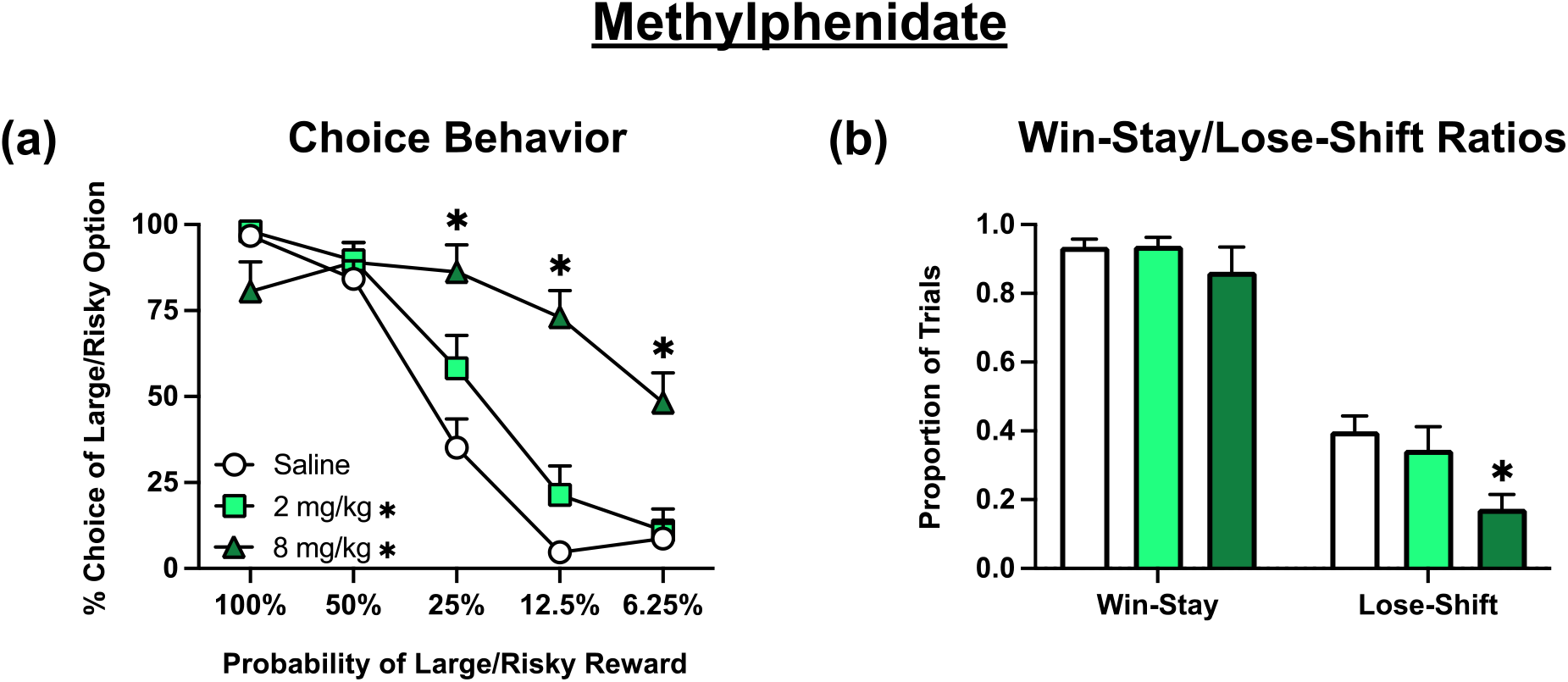
Effects of methylphenidate (n = 15) during probabilistic discounting. All conventions are the same as Fig. 1. (**a)** Choice Behavior: * on symbol key denotes p < 0.05 for an overall treatment effect of each dose versus vehicle. * on graph denotes p < 0.001 for the 8 mg/kg doses versus vehicle at a specific block analyzed by Dunnett’s multiple comparisons tests. (**b)** Win-Stay/Lose-Shift Ratios: * on graph denotes p < 0.001 versus vehicle analyzed by Dunnett’s multiple comparisons tests following one-way ANOVA analysis. Symbols and bars represent mean ± SEM.

Analysis of win-stay/lose-shift data (**Fig. 2b**) revealed a significant main effect of both feedback [F (1.000, 14.00) = 93.84, p < 0.0001], and treatment [F (1.682, 23.55) = 12.58, p = 0.0004], but did not yield a feedback x treatment interaction [F (1.511, 21.16) = 1.203, p = 0.3078]. However, inspection of Fig. 2b suggest that the effect of treatment was driven primarily by a reduction in lose-shift behavior. This was confirmed by two, exploratory one-way ANOVAs that revealed a lack of effect of treatment for win-stay behavior [F (1.258, 17.62) = 0.8829, p = 0.3845], whereas analysis of lose-shift data did reveal a main effect of treatment [F (1.766, 24.73) = 9.559; p = 0.0012]. Dunnett’s multiple comparisons determined that the 8 mg/kg dose of MPH significantly decreased lose-shift tendencies (p = 0.0007) compared to vehicle treatments Analysis of choice latency data and trial omissions (**Table 1**) failed to reveal a main effect of block [F (1.965, 27.51) = 0.2085, p = 0.8093; F (2.242, 31.39) = 0.7375, p = 0.5009, respectively], but no main effects of treatment [F (1.798, 25.17) = 1.706, p = 0.2036; F (1.435, 20.09) = 1.688, p = 1.688, respectively] or block x treatment interaction [F (2.750, 37.81) = 0.8910, p = 0.4471; F (2.338, 32.73) = 1.743, p = 0.1865, respectively].

Analysis of magazine latency data (**Table 1**) revealed a significant main effect of block [F (1.476, 20.66) = 7.658, p = 0.0060], but did not yield a significant main effect of treatment [F (1.779, 24.90) = 0.5905, p = 0.5426] or a block x treatment interaction [F (2.992, 38.53) = 1.936, p = 0.1401].

Overall, these results show that MPH, another psychostimulant with cognition-enhancing properties, also increases risky choice preference in this assay.

### Effects of SK609

In contrast to the two other psychostimulant drugs tested, analysis of the effects of SK609 on choice (**Fig. 3a**) did not yield a significant main effect of treatment [F (1.000, 11.00) = 1.430, p = 0.2569] or a block x treatment interaction [F (2.326, 25.58) = 0.01624, p = 0.9909].

**Fig. 3.**
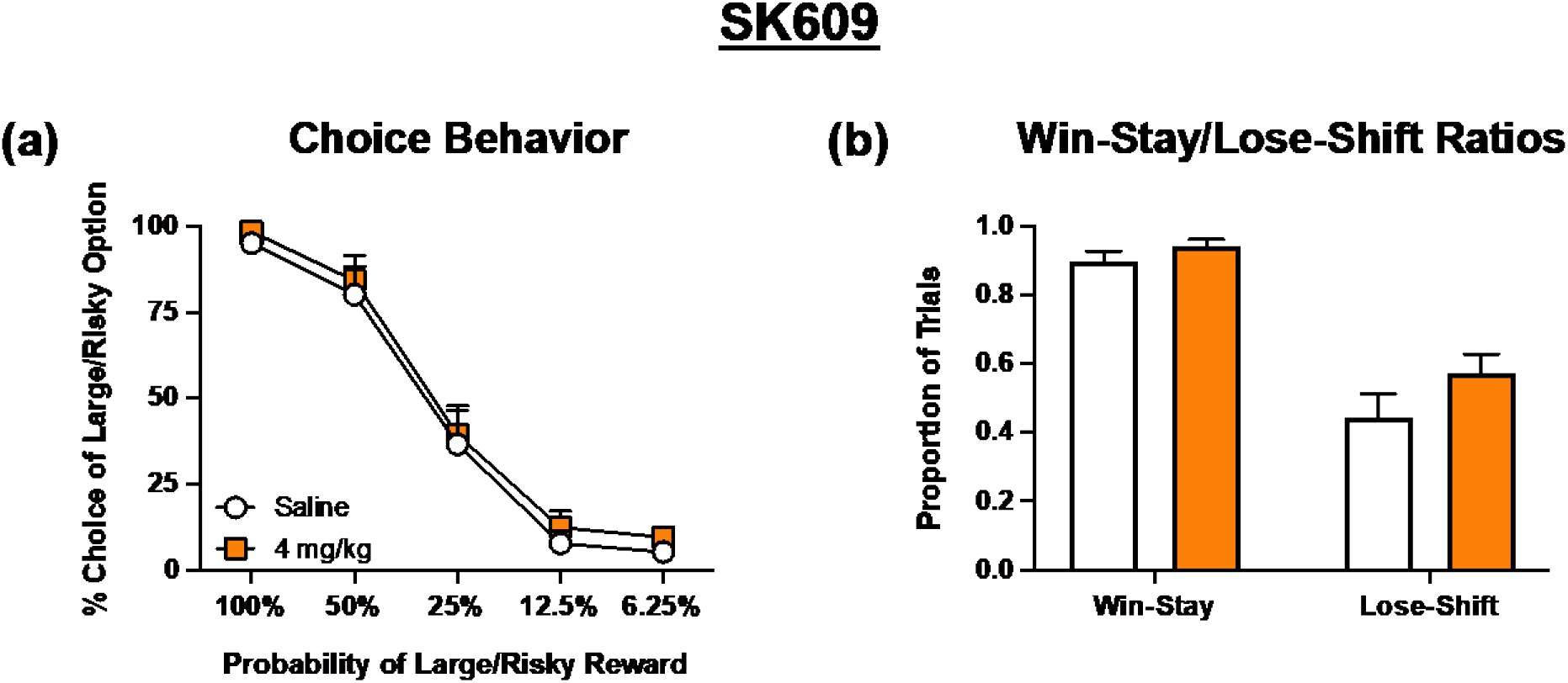
Effects of SK609 (n = 12) during probabilistic discounting. All conventions are the same as Fig. 1. (**a)** Choice Behavior: no specific effects observed. (**b)** Win-Stay/Lose-Shift Ratios: SK609 trended towards a treatment effect at p = 0.0693. Symbols and bars represent mean ± SEM.

Analysis of win-stay/lose-shift data (**Fig. 3b**) revealed a significant main effect of feedback [F (1.000, 11.00) = 53.69, p < 0.0001] and a strong trend towards a significant main effect of treatment [F (1.000, 11.00) = 4.049, p = 0.0693], suggesting that this compound subtly increased both win-stay and lose shift behavior. There was no significant feedback x treatment interaction [F (1.000, 11.00) = 1.247, p = 0.2879] observed.

Analysis of choice latency data (**Table 1**) revealed a significant main effect of block [F (1.373, 15.10) = 5.503, p = 0.0246], but did not yield a significant main effect of treatment [F (1.000, 11.00) = 0.01082, p = 0.9190] or a block x treatment interaction [F (1.982, 21.80) = 0.2375, p = 0.7887].

Analysis of magazine latency data and trial omissions (**Table 1**) failed to reveal a main effect of block [F (1.061, 11.67) = 0.9101, p = 0.3657; F (1.373, 15.10) = 3.283, p = 0.0797; respectively], a main effect of treatment [F (1.000, 11.00) = 2.460, p = 0.1450; F (1.000, 11.00) = 1.093, p = 0.3182, respectively] or a block x treatment interaction [F (1.035, 11.38) = 2.189, p = 0.1661; F (1.674, 18.42) = 0.4256, p = 0.6247; respectively].

Overall, these results show that SK609 does not increase risky choice behavior at its cognitive enhancing dose as represented by the lack of change in the shape of the probabilistic discounting curve.

## Discussion

The main objective of the present study was to evaluate three agents with cognitive-enhancing properties for their potential to alter risk/reward decisions, examining their potential side effect profile of increasing risky choice. While all three drugs have been shown to improve selected dimensions of PFC-mediated cognitive function, this data shows the effects of SK609 can be dissociated from the psychostimulant class of drugs using the PDT. AMPH and MPH increased risky choice behavior at their cognition-improving doses, whereas SK609 did not.

As has been reported previously, AMPH induced a marked increase in risky choice when reward probabilities were initially high and then diminished over a session, and this was associated with a reduced sensitivity to non-rewarded choices. Note that the increase in risky choice described here is unlikely to reflect an overall increase in risk preference, as previous studies have shown that AMPH can actually reduce risky choice on this task when reward probabilities are initially low and increase over a session (St. Onge et al., 2010). Thus, AMPH appears to induce a “inflexible” phenotype, making it more difficult for animals to shift decision bias away from their preferred option at the start of the session as reward probabilities change. In this instance, rats were less affected by non-rewarded choices as reward odds decreased and persisted in selecting the risky option even when it had less utility. Thus, although AMPH can enhance certain forms of cognition and executive functioning, it can exert somewhat deleterious effects on flexible decision making guided by choice/outcome history.

While it is clear that stimulants and non-stimulants have differing potentials to increase risk-taking behavior, there are also differences between MPH and AMPH. Interestingly, the effects of MPH on this form of probabilistic discounting had not yet been evaluated. MPH induced a moderate increase in risky choice at its cognitive-enhancing dose; however, it did not alter feedback sensitivity. AMPH not only increased risky choice at all doses, but these effects were driven by reduced sensitivity to non-rewarded risky choices, a behavior observed only at the highest dose (8 mg/kg) of MPH. These results are similar to those recently reported using a dependent schedule variant of a probability discounting task, where both drugs increased risky choice and decreased sensitivity to probabilistic reinforcement, but only the highest dose of MPH significantly altered the latter (Yates et al., 2020). These differential effects may be due to the different mechanisms by which MPH and AMPH enhance catecholamine transmission. MPH and AMPH both block the DA and NE transporters; however, AMPH also enhances catecholamine release through reverse transport (Leyton et al., 2002; Heal et al., 2013; Faraone, 2018). In combination, these mechanisms may lead to supranormal increases in catecholamine activity and excessive receptor stimulation, resulting in a secondary effect on feedback sensitivity. It may be that the high 8 mg/kg dose of MPH mimics these effects whereas the low 2 mg/kg dose of MPH may not be sufficient to alter lose-shift behavior. Nevertheless, these findings indicate that while psychostimulants can promote risky behavior, not all aspects of risk/reward decision making are affected in the same way. MPH may be a more favorable option over AMPH when considering treatment options for improving cognitive performance while limiting side-effect symptoms.

Non-stimulant therapies have become a favorable alternative to stimulant treatments because of their minimal potential for substance misuse and fewer side effects (Pliszka, 2003). Here, we observed that SK609, which selectively blocks NE transporters without affinity for DA transporters, did not alter choice preference. That said, this compound did induce a trend towards enhanced sensitivity to both rewarded and non-rewarded feedback, which may have offset overall changes in risky choice. Atomoxetine (ATX), another selective NE reuptake blocker, is a non-stimulant agent used for treating ADHD symptoms (Michelson et al., 2003; Michelson et al., 2001; Durell et al., 2013) and is known for improving rodent cognitive performance and impulse control in other tasks (Cain et al., 2011; Robinson et al., 2008; Chernoff et al., 2021; Chernoff et al., 2023; Navarra et al., 2008). Surprisingly, a low dose of ATX (0.3 mg/kg) has been shown to increase risk preference in the PDT in the more advantageous, higher probability blocks, whereas slightly higher doses (1 and 3 mg/kg) did not (Montes et al., 2015). However, ATX either had no effect or induced optimal choice strategies in similar rodent gambling tasks of risky choice preference (Baarendse et al., 2013; Chernoff et al., 2023; Chernoff et al., 2021). Thus, the selective increases in risky choice induced by NET blockade during the highest large reward probability blocks using an ascending limb protocol may reflect an improvement in the ability to optimize behavioral strategy to maximize gains. Notably, while all three doses of ATX previously profiled in the PDT are considered clinically relevant, the lowest dose of ATX has greater selectivity for the NE transporter compared to the higher doses, which also block other monoamine reuptake (Bymaster et al., 2002). In addition to NE transporter blockade, SK609 selectively activates DA D3 over all other DA receptors, a mechanism previously shown alone to reduce risky choice preference in the PDT (St Onge and Floresco, 2009). It is highly plausible that any increases in risky choice preference that could have been driven by NE transporter blockade were offset by this DA D3 stimulation.

Interestingly, no changes in choice or magazine latencies were observed across all treatment conditions.
Although AMPH and high doses of MPH decreased lose-shift tendencies, this reduced sensitivity to non-rewarded outcomes does not appear to promote greater or lesser deliberation of subsequent decisions. The lack of altered choice latencies was also surprising given the potential hyperactivity effects that are typically observed with higher doses of these drugs and associated with increased DA activity in subcortical regions of the brain (Arnsten, 2006; Berridge et al., 2006; Devilbiss and Berridge, 2008; Schmeichel and Berridge, 2013). AMPH users have been shown to exhibit slower deliberation times in comparison to control patients within a clinical decision-making task (Rogers et al., 1999). Therefore, it is possible that intermittent dosing of AMPH and MPH is not sufficient to cause significant changes in choice latencies but continuous/repeated long-term exposure to stimulant agents can lead to alterations in the rate in which cost/benefit contingencies are effectively evaluated. Given that SK609 did not alter choice preference or feedback sensitivity, it is unsurprising that choice and magazine latencies were also unaffected. Taken together, these results suggest that stimulant and non-stimulant agents do not significantly affect the rate at which cost/benefit contingencies are evaluated and the desire to collect one’s reward.

Together, these results highlight the roles of NE transporter blockade and selective D3 activation in pro-cognitive action without side effect liability. In marked contrast to the disruptive effects of AMPH and MPH on risky choice, SK609 did not alter risky choice or any other behavioral measures. The absence of DA transporter blockade and non-selective dopaminergic elevation are beneficial properties of SK609 that differentiate it from traditional pro-cognitive psychostimulants. These data show that SK609, which produces favorable effects on multiple domains of PFC-mediated cognitive function (Schneider et al., 2021; Marshall et al., 2019), does not induce psychostimulant-like side effect liability at its cognitive-enhancing dose when assessed with the PDT.

## Conclusion

Overall, our results highlight a potential side-effect of stimulant compounds that should be considered when prescribing treatment options for patients with PFC-mediated cognitive dysfunctions. SK609 offers an alternative to traditional psychostimulants for improving cognition without promoting risk taking behavior.

## Conflict of interest statement

On behalf of all authors, the corresponding author states that there are no conflicts of interest.

## Acknowledgements of Funding

This study was supported by the New Jersey Commission on Brain Injury Research grants: CBIR20PIL004 to R.L.N. and CBIR19IRG025 to B.D.W., the United States Department of Defense Traumatic Brain Injury and Physiological Health Research Program grant W81XWH-22-1-0616 to R.L.N. and W81XWH-22-1-0618 to B.D.W., and the Osteopathic Heritage Foundation for Primary Care Research Award to R.L.N.

## Notes

### Competing Interest Statement

The authors have declared no competing interest.

